# Establishment of a publicly available core genome multilocus sequence typing scheme for *Clostridium perfringens*

**DOI:** 10.1101/2021.04.20.440723

**Authors:** Mostafa Y. Abdel-Glil, Prasad Thomas, Jörg Linde, Keith A. Jolley, Dag Harmsen, Lothar H. Wieler, Heinrich Neubauer, Christian Seyboldt

## Abstract

*Clostridium perfringens* is a spore-forming anaerobic pathogen responsible for a variety of histotoxic and intestinal infections in humans and animals. High-resolution genotyping aiming to identify bacteria at strain level has become increasingly important in modern microbiology to understand pathogen transmission pathways and to tackle infection sources. This study aimed at establishing a publicly available genome-wide multilocus sequence-typing scheme for *C. perfringens*. 1,431 highly conserved core genes (1.34 megabases; 50% of the reference genome genes) were indexed for a core-genome-based MLST scheme for *C. perfringens*. As an example, we applied the scheme to 87 poultry and 73 non-poultry strains (total=160). The genotyping results of the 160 genomes were congruent in terms of resolution and tree topology between allele-based and single-nucleotide-polymorphism-based core-genome typing. For the analysis of poultry strains of *C. perfringens* concerning the country of isolation, NetB-toxin gene carriage and clinical disease, we used 60 allelic differences as a clustering threshold. The results showed that poultry strains from a single country formed a cluster (*n=*17 clusters including 46 strains). Two clusters included six strains from four different countries. These strains were *netB*-positive, as were seven strains from Denmark and two strains from Finland, possibly indicating common sources of *netB*-positive strains. In terms of clinical presentation, different clusters of strains were associated with cases of suspected necrotic enteritis. Strains from sick birds grouped with strains from healthy birds or meat samples showing that potentially virulent strains are widespread and that host-related factors contribute significantly to NE. In summary, a publicly available scheme and an allele nomenclature database for genomic typing of *C. perfringens* has been established and can be used for broad-based and standardised epidemiological studies.

## Introduction

*Clostridium perfringens* is a Gram-positive anaerobic bacterium that is widely distributed in the soil and faeces of humans and animals (1). *C. perfringens* produces resistant spores that allow the bacterium to survive in harsh environments and play a central role in the epidemiology of *C. perfringens* diseases (2). This bacterium produces a wide array of extracellular toxins and enzymes. Based on the presence of six typing toxins (α, β, ε, ι, CPE and NetB), *C. perfringens* is classified into seven toxinotypes (A to G) (3). The toxins used for typing are plasmid-encoded except α-toxin that is chromosomally encoded and CPE where the gene can be located on a chromosome or a plasmid (4). Of the seven toxinotypes, *C. perfringens* type A is the most common and is widely isolated from healthy individuals and the environment. The diseases caused by type A strains are diverse including traumatic gas gangrene in several mammalian species for which α-toxin and θ-toxin are important in disease progression (5). Type A also causes a variety of enteric infections in domestic animals e.g. yellow lamb disease in sheep and necrohaemorrhagic enteritis in calves (1, 6). In contrast to type A, toxinotypes B to G are associated with the incidence of certain diseases in specific host(s). For example, *C. perfringens* type B strains cause fatal haemorrhagic dysentery in lambs (1, 4) while type C strains cause necrotic enteritis and enterotoxaemia in lambs, piglets, calves and foals. Type C strains that produce CPE and β toxin are associated with foodborne illness in humans known as Pigbel or Darmbrand. *C. perfringens* type D is implicated in enterotoxaemia in sheep and goats (4). Type E strains are occasionally associated with calf enterotoxaemia and haemorrhagic enteritis (4). Type F strains produce CPE that is particularly important in *C. perfringens* food poisoning as well as non-foodborne gastroenteritis in humans. *C. perfringens* type G describes the *netB*-positive strains that cause necrotic enteritis (NE) in poultry (7).

Bacterial strain typing is essential for outbreak investigations, epidemiological surveillance and evaluation of control measures (8). Multilocus sequence typing (MLST) has been widely used for microbial genotyping. MLST provides portable data easy for comparison among different laboratories (9). Owing to the significantly reduced costs of DNA sequencing over time, MLST has been extended to involve many hundreds of genes with the so-called core genome MLST (cgMLST) and whole-genome MLST (wgMLST) providing high resolution for optimal pathogen typing (10).

In *C. perfringens*, a cgMLST system was firstly utilised to describe the clonal relationship between *netF*-positive strains from enteritis cases in foals and dogs (11). 1,349 genes were used to type 47 *C. perfringens* strains (11). The results showed that 32 *netF*-strains were classified into two clusters including 26 and 6 strains, respectively (11). The scheme used in this previous study has not been validated and is not currently available for subsequent typing. This underscores the need to standardise an accessible, genome-wide typing approach for a uniform and replicable characterisation of *C. perfringens*. The aim of the current study was therefore to develop a cgMLST scheme for *C. perfringens*. The scheme was applied to study 160 *C. perfringens* genomes: 87 of poultry origin and 73 of non-poultry origins.

## Methods

### Bacterial strains and whole-genome sequence data retrieval

A total of 160 *C. perfringens* genome sequence were involved in this study (Table S1, Table 1). The genomes of 80 diverse *C. perfringens* strains available at NCBI were used to infer the species by phylogenetic analysis and to select representative genomes of *C. perfringens*. These 80 genomes were additionally used for initial evaluation and refinement of the cgMLST scheme via removing frequently untypeable genes. The origin of these strains in respect to isolation time, geography and host is available at Table S1.

**Table 1:**
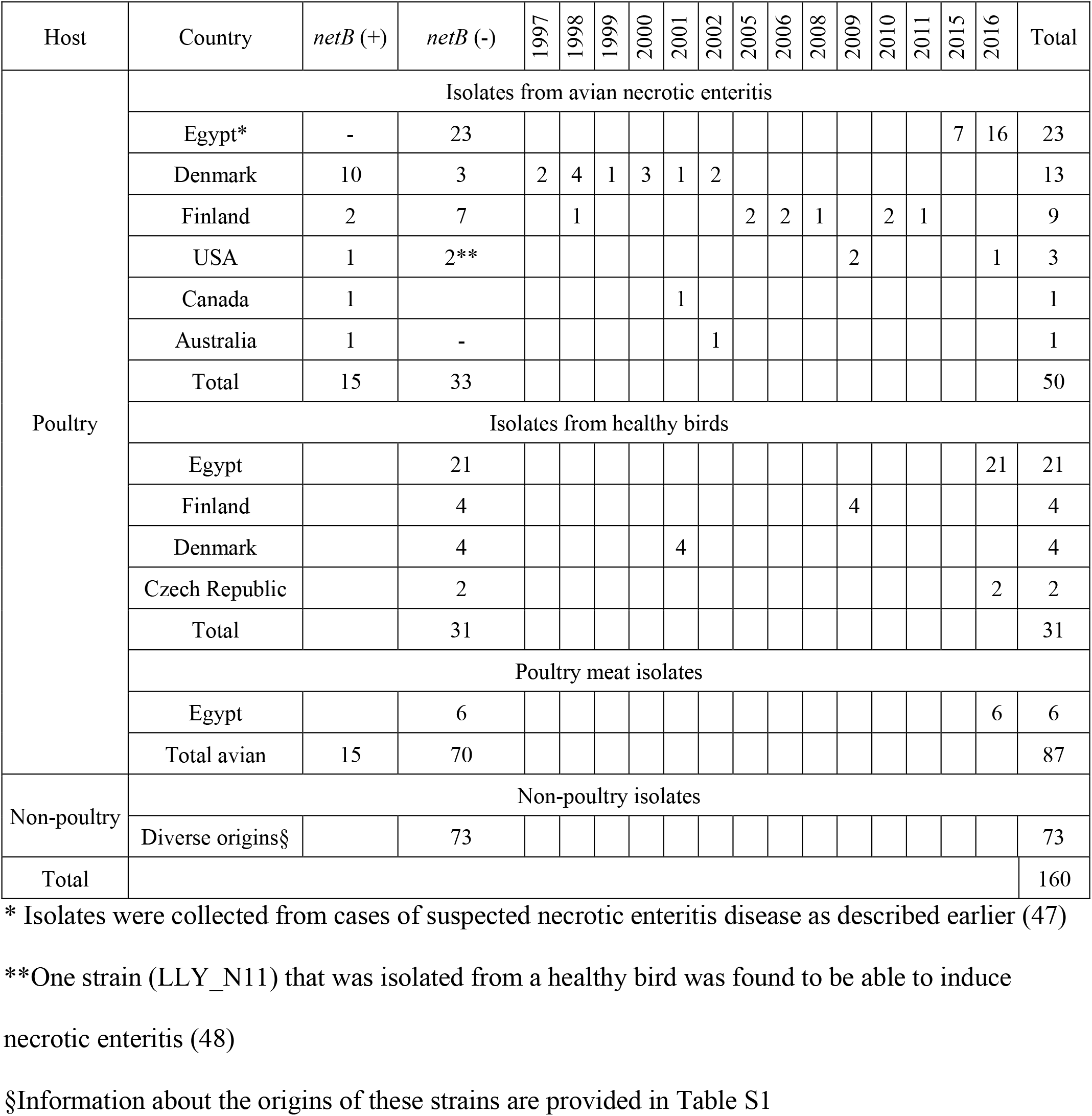
Overview of all *Clostridium perfringens* strains included in the core genome multilocus sequence typing scheme definition and validation.

To apply the scheme to poultry strains of *C. perfringens*, we sequenced 50 *C. perfringens* strains from Egypt (Table S1). These strains comprise (i) 23 strains from cases suspected for NE in 14 farms including the disease pictures clinical (*n=*17) and subclinical (*n=*6), (ii) 21 strains collected from clinically asymptomatic birds at slaughterhouses, and (iii) six strains from retail chicken meat parts. In addition, publicly available raw sequencing data (Illumina MiSeq) of 30 poultry strains from Finland and Denmark were included (12) (Table S1). The strains span a time period between 1997 and 2010 and include (i) 17 strains derived from healthy (*n=*4) and NE infected (*n=*13) chickens in Denmark and (ii) 13 strains obtained from healthy (*n=*4) and NE infected (*n=*9) turkeys in Finland (12).

### Extraction of genomic DNA and WGS of *C. perfringens* from poultry from Egypt

*C. perfringens* strains (*n=*50 strains) were cultured on Schaedler Agar with Sheep Blood (CM0437 Oxoid, Germany) incubated overnight at 37°C under anaerobiosis. 1-5 fresh colonies were inoculated in 13mL Selzer broth (13) (tryptone 30g, beef extract 20g, glucose 4g, L-cysteine hydrochloride 1g in 1000mL H2O, pH 7.2) and incubated under anaerobiosis for 2.5-3hrs. 1.5mL of the culture was pelleted (13500 rpm for 10 min) and used for the extraction of DNA with DNeasy Blood and Tissue Kit (Qiagen, Germany). For eluting nucleic acids, Buffer EB (10 mM Tris-Cl, pH 8.5, Qiagen, Germany) was used. The quality of DNA was assessed as follows: (i) the concentration and the purity (absorbance ratio 260/230 and 260/280) of the DNA were measured using a NanoDrop spectrometer (Thermo Fisher Scientific, USA). (ii) The integrity of the DNA was analysed by agarose gel electrophoresis. (iii) A Qubit 2.0 fluorometer (Life Technologies, Germany) was used to calculate the concentration of the double-stranded DNA (dsDNA). The Qubit dsDNA BR Assay Kit (Q32851, Life Technologies, Germany) was used for DNA quantification according to the manufacturers’ instructions. Whole-genome sequencing was performed by GATC Biotech (Konstanz, Germany) with paired-end libraries on an Illumina HiSeq platform (Illumina, USA) generating reads of 151bp in length.

### Genome *de novo* assembly and annotation

First, the quality of the paired-end Illumina reads was checked using FastQC (Babraham Bioinformatics, Cambridge) and trimmed using Sickle v1.33 (https://github.com/najoshi/sickle). Genome assembly was performed using SPAdes v3.9.1 (14) with mismatch correction settings. Generated contigs were filtered for a kmer coverage threshold of 3-fold and a minimum contig length of 500bp using an in-house script. Genome annotation was performed using Prokka v1.11 (15).

### Core genome multilocus sequence typing scheme definition

Fig. 1 visualises the complete workflow to define and evaluate the core genome MLST targets. The genome sequence of the type strain ATCC 13124 (accession NC_008261.1) was used as a reference (16). The following filters were then applied using SeqSphere+ v7.1.0: (i) a “minimum length filter” to discard genes of length less than 50 bp, (ii) a “start codon filter” and “stop codon filter” to discard genes that lack one start- and /or one-stop codon, (iii) a “homologous gene filter” for discarding genes that occur in multiple copies at a sequence identity of at least 90% over a region of 100bp or more, and (iv) a “gene overlap filter” that if two genes overlap by more than 4bp length, the shorter gene will be discarded.

**Figure 1:**
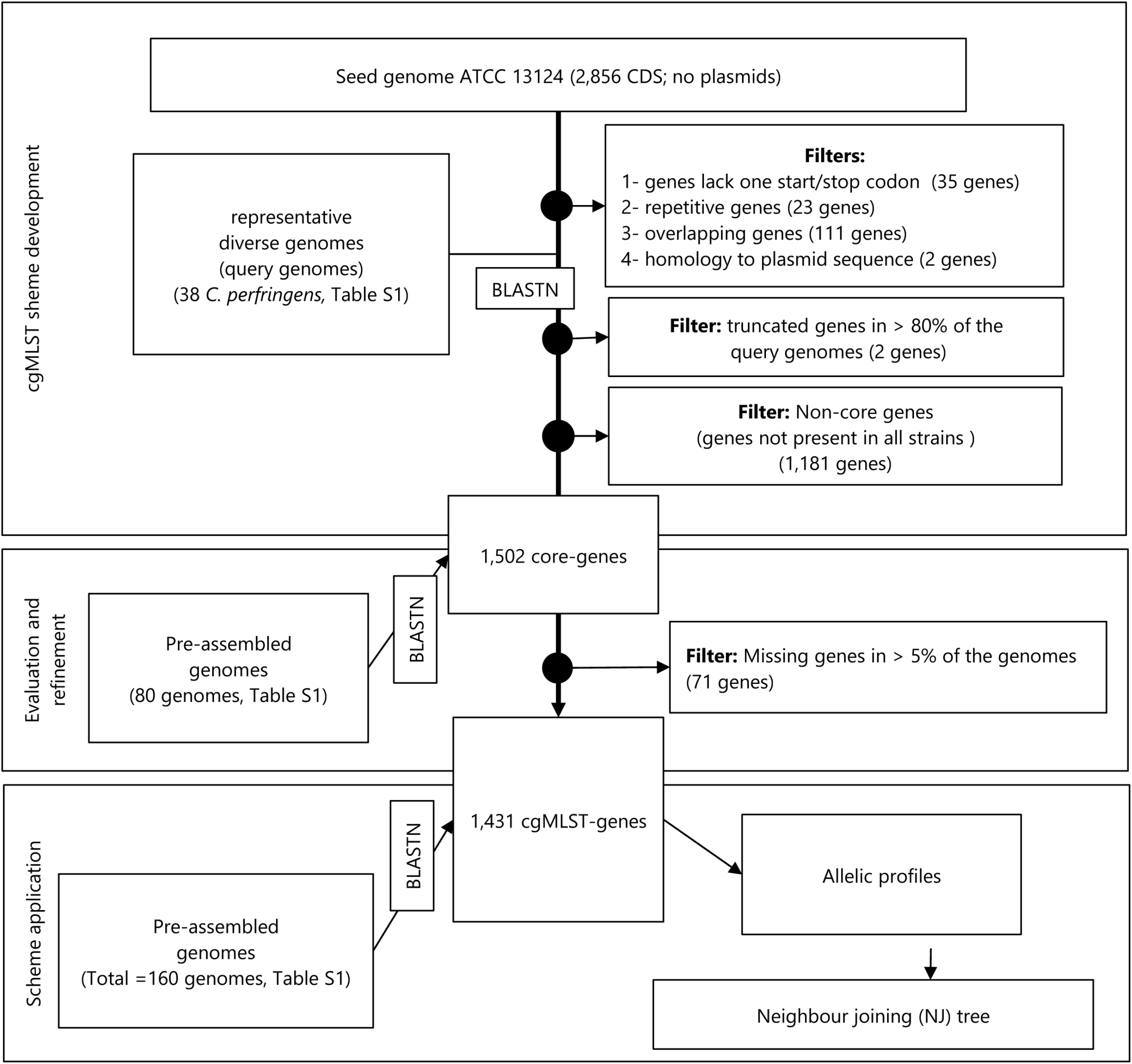
Scheme of the workflow to define and evaluate the core genome MLST targets.

To determine the species core genes, the retained genes from the reference strain were compared with BLAST against 38 *C. perfringens* genomes (also called query genomes). These genomes comprised 26 closed genomes and 12 draft assemblies that represent the five major phylogroups of *C. perfringens*, as recently described (17). Based on the size and sequence divergence for each phylogroup, the number of genomes selected was as follows: 12 strains for phylogroup I, 23 strains for phylogroup III and one strain each from phylogroups II, IV and V (Table S1, Figure S1). By default, the program SeqSphere excludes genes with internal stop codons in more than 80% of the query genomes. BLAST v2.2.12 was used at 90% identity and 100% coverage. BLAST options included match reward 1, word size 11, gap open costs 5, mismatch penalty -1 and gap extension cost 2. To further exclude genes with homology to plasmids, we constructed a database of *C. perfringens* plasmid sequences (*n*=90 plasmids) available in GenBank or recently assembled based on PacBio sequences (17). As a result, we filtered out genes with a BLAST match to any of the plasmid sequences at 90% identity.

Next, we aimed to evaluate the core genes to keep only the complete genes that are most likely stable in *C. perfringens*. For this purpose, we used a set of 80 genomes including the above-mentioned 38 genomes (Table S1). The genomes in this data set are publicly available and are geographically and temporally divergent. These genomes were processed using the SeqSphere+ v7.1.0 searching for each of the defined core genes in the assembled genomes. Parameters included 90% identity and 99% overlap to reference sequence. Only complete genes were assigned to allele numbers i.e. alleles were not assigned to genes with frameshifts, in-frame stop codons or carry non-GATC characters. In-frame multiple insertions or deletions (indels) were allowed up to three codons per gene relative to the reference genes. Uncalled genes due to gene absence or failed allele assignments in more than 5% from the initial data set were excluded. This final set of the filtered core genes served as cgMLST targets (Table S2). The Pairwise Homoplasy Index (PHI) (18) was calculated using PHIPack as a statistical method for detecting the presence of recombination in the final cgMLST genes.

### Application of the core genome multilocus sequence typing scheme

According to the final set of cgMLST targets, the poultry-specific *C. perfringens* strains were investigated concerning clinical disease, geography and NetB toxin gene carriage. For that, an additional set of 80 poultry *C. perfringens* strains was used (Table S1) including 50 strains from Egypt and 30 strains from Denmark and Finland. All strains (*n=*160) were assigned allele profiles according to the final cgMLST targets.

### Whole genome-based single nucleotide polymorphism (SNP) detection

The RedDog pipeline v1beta.10.3 (https://github.com/katholt/reddog) was used to detect SNP variations in the whole genome of 160 *C. perfringens* strains. Of these, Illumina reads were not available for 61 genomes that were derived from GenBank. Therefore, simulated reads (one million 100bp reads) were generated for the 61 genomes using the SAMtools v1.4-14 (19). The genome of strain ATCC 13124 (16) served as reference for SNP calling. Illumina reads as well as simulated reads were mapped to the reference sequence using Bowtie2 v2.3.0 with a sensitive-local algorithm and a maximum insert length of 2000 (set via -x option) (20). In each genome, SAMtools v1.4-14 (19) was used to call SNPs with Phred quality score ≥ 30 and read depth ≥5x. Ambiguous SNPs including heterozygous allele calls were discarded. In addition, repeat sequences and prophage regions were identified in the reference strain using nucmer (21) and PHASTER (22), respectively. SNPs located in these regions were excluded. Whole-genome SNP sites that were present in the coding and non-coding sequences and conserved in 99% of the genomes were selected. RAxML v8.2.10 was used to generate an ML phylogenetic tree using the general time-reversible (GTR)-gamma model and 100 bootstrap replicates (23).

In order to generate a recombination filtered SNP tree, Gubbins v.2.2.1 was used to identify and mask putative recombination regions from the genome alignment (24). As input to Gubbins, a pseudomolecule was constructed for each strain consisting of the reference sequence (ATCC 13124) but updated with sample-specific SNPs using snpTable2GenomeAlignment.py available at (https://github.com/katholt/reddog).

### Comparison between cgMLST and whole genome SNP typing

To visualise the topological concordance between the cgMLST neighbor joining (NJ) tree and the SNP-based maximum likelihood tree, the tanglegram algorithm was applied (25) using Dendroscope v.3.2.1027 (26). The tanglegram method allows for a direct comparison between both trees via plotting them side by side and matching the corresponding taxa. Before generating the tanglegrams, both trees were rooted with strain CBA7123 (accession NZ_AP017630.1).

To compare the discrimination levels and assess the congruence of the typing results using cgMLST as well as SNP typing methods, the Simpson’s diversity index (SDI) (27) and the adjusted Wallace index of concordance (28) as calculated using the Comparing Partitions tool (29) were applied. The SDI (27) is a quantitative measure of diversity which denotes the probability that two individuals randomly selected from a sample will be classified into two different types. SDI values range between zero and one where one represents the maximum diversity in the sample. The adjusted Wallace’s coefficient (28) measures the probability that two individuals classified together using method A will also have the same classification type by method B.

### Comparison of cgMLST typing to classical MLST typing methods

To compare the resolution obtained by cgMLST to the resolution of classical MLSTs, the MLST loci of classical schemes were extracted *in silico* from the WGS data of the 160 strains using BLASTN (30). Three classical MLST schemes previously described for *C. perfringens* were involved (31-33). The MLST genes described by Deguchi et al., (2009) (32) include *groEL, gyrB, nadA, pgk, sigK, sodF, plc* and *colA*, the genes described by Jost et al., (2006) (31) include *cpa, ddlA, dut, glpK, gmk, recA, sod* and *tpiA* while the MLST genes described by Hibberd et al., (2011) (33) include *dut, ddl, sod, tpi, dnaK, glpK, gmk, gyrA, plc, recA* and *groEL*. For each MLST locus, the extracted sequences from the 160 genomes were aligned using MAFFT (34) and then assigned allele numbers and sequence types using MLSTest (35). Simpson’s Index of Diversity (27) and adjusted Wallace coefficients (28) were calculated using the Comparing Partitions tool (29) to assess the discriminatory power and to evaluate the congruence of the typing results between cgMLST and classical MLSTs methods. The topological congruence of the NJ trees from cgMLST and classical MLSTs was visualised with the tanglegram algorithm (25) using Dendroscope v.3.2.1027 (26). All trees were rooted with strain CBA7123 before creating tanglegrams.

### Data Availability

The generated raw sequencing data are deposited at NCBI under the BioProject PRJNA718992.

## Results

### Development of a cgMLST scheme

The reference genome of strain ATCC 13124 carries 2,856 CDSs, of which 171 genes were removed after applying the basic filters of SeqSphere+ v7.1.0 (Fig. 1). 1,502 genes were strictly present in the 38 query genomes, representing core genes. These genes were evaluated to select the most suitable genes for cgMLST typing. Therefore, allele typing was performed using 80 *C. perfringens* strains of diverse origins (Table S1). Allele numbers were assigned only to the complete genes without ambiguities. This resulted in the exclusion of a further 71 genes to which no allele numbers were assigned in >5% of the investigated genomes (Fig. 1, Table S3). The final cgMLST genes thus comprised 1,431 genes (1.34Mbp), corresponding to 41.2% of the reference genome size (Table S2). The average GC content of these genes was 29.8% with lengths between 84 and 4,350bp (average=938.2bp) (Fig. 2).

**Figure 2:**
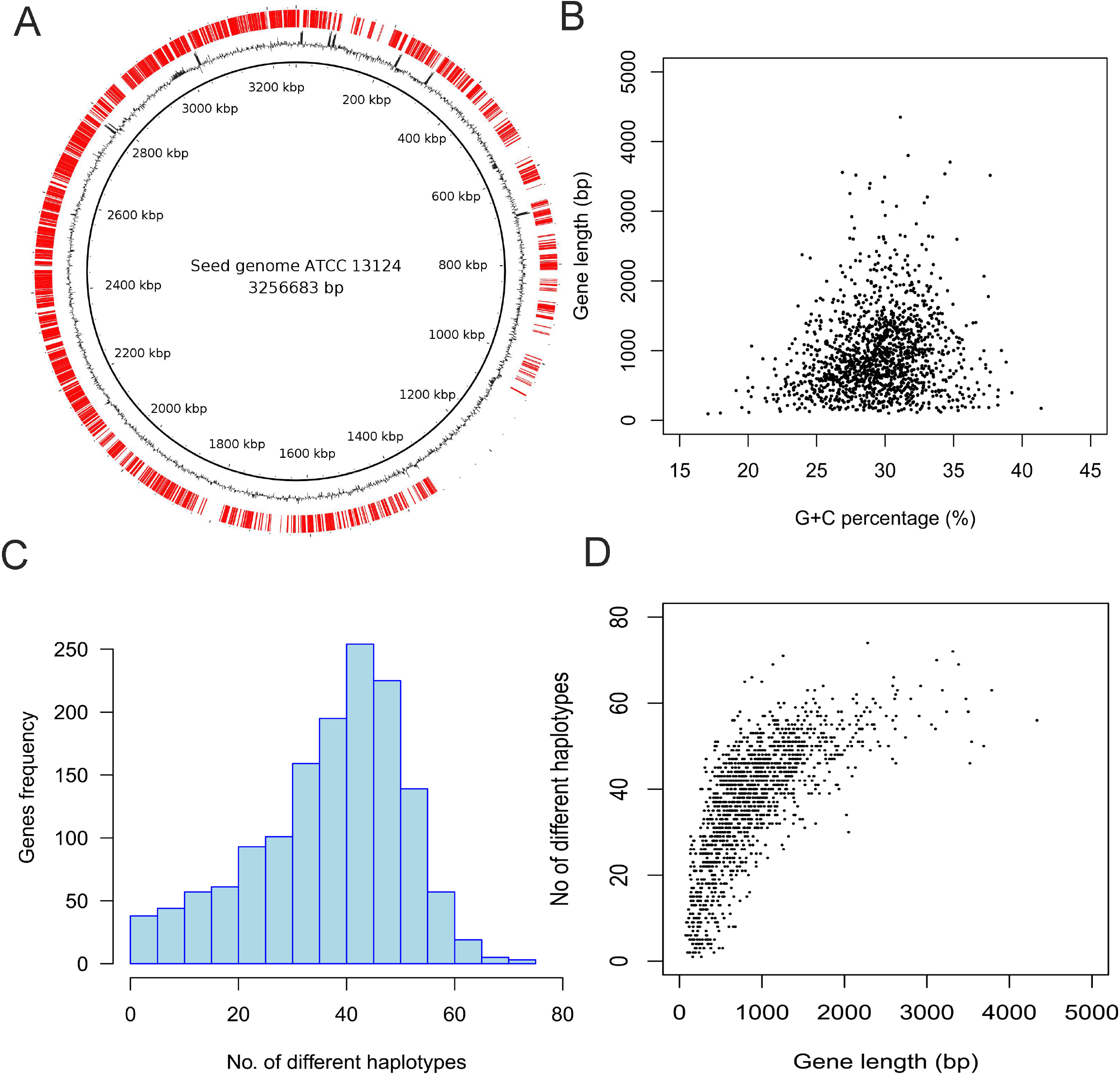
Characteristics of the 1,431 core genome MLST genes. (A) The distribution of the cgMLST targets (red, outer circle) across the *Clostridium perfringens* ATCC 13124 reference genome. Inner circles represent the G+C content and genomic positions, respectively. (B) Plot showing the lengths and G+C content of the cgMLST targets. (C) Plot showing the frequency of different allelic variants observed for each of the cgMLST targets. (D) Plot showing a direct association between the length of cgMLST target genes and the number of variants detected for each gene.

### Application and performance of cgMLST

For scheme application, an additional genome set of 80 poultry strains of *C. perfringens* was used (Table S1). Genome assembly resulted in a high-quality draft genome sequence for most strains with an average contig number of 79±66 and N50 of 543.2±521.1Kbp (Table S4). The 50 strains from Egypt yielded an average sequencing depth of 792.2±277-fold while the 30 strains from Finland and Denmark yielded an average sequencing depth of 103.7±49-fold (Table S4).

On average 99.5% of the cgMLST genes were identified and assigned allele numbers. The non-typeable genes ranged between 0 and 71 (median=5) per genome totalling 568 genes. The absence of a BLAST match and/or the presence of internal stop codons or ambiguous nucleotides are the reasons for the failure of allele assignment. On average 2.58±5.9 genes per genome had no BLAST match (*n*=283 genes). Three genes (CPF_RS09035, CPF_RS07255 and CPF_RS09515) had no BLAST match in >10 genomes. Additionally, the genome of the strains T43, T34, JGS1721, C7 and C3 exhibited the highest number of non-detected genes with BLAST in which 47, 40, 27, 26 and 20 genes were not detected, respectively. The assembled genome data of these strains are characterised by high contig numbers and low N50 values, indicating excessive fragmentation of these genomes (Table S4).

Errors due to frameshifts, internal stop codons or ambiguous nucleotides resulted in the failed allele assignments to 365 target genes in at least one genome, with an average of 4.8 ± 5.7 (range 0 to 40) genes per strain. Of these, six loci (CPF_RS05860, CPF_RS04475, CPF_RS10790, CPF_RS13045, CPF_RS07050 and CPF_RS13485) were not assigned allele numbers in more than ten genomes. It is noteworthy to mention that failed targets due to frameshifts and internal stop codons were reported at high frequency in the closed genome of strain 13 in which alleles for 29 targets were not assigned, 26 thereof due to indel-associated frameshifts and the other three genes due to substitution that led to an internal stop codon. In total, indel mutations were observed in a sum of 298 genes, ranging from 1 to 13 nucleotide differences per gene relative to the cgMLST targets (Table S2).

The average number of alleles reported for each cgMLST gene including the entire data set was 37.2±13.8 alleles (range 1 to 74) (Fig. 2, Table S2). The cgMLST system assigned nearly every isolate as a distinct sequence type, as the number of distinct allelic profiles was 156 for the 160 genomes. The SDI for the cgMLST was 1.0 (95% confidence interval [0.999-1.0]). Of interest, homologous recombination was found to affect nearly half of the loci as determined by the PHI statistic test (883 out of 1,431 loci with *P* value less than 0.05) (Table S2).

### Definition of cgMLST cluster types

Similar to clonal complexes for classical MLST, cgMLST profiles can be grouped into Cluster Types (CT) defining a group of very similar cgMLST profiles differing by up to a certain number of alleles defined as CT threshold. In this study, we used a CT threshold of 60 allelic differences because it showed a good concordance with a previous study that grouped the *netF*-positive strains into two clusters (11). We also calculated the allelic distances between each pair of genomes and found that the pairwise allelic distances were less than 60 for more than half of the genomes (84 strains) accounting for ∼3% of the overall pairwise allelic distances (∼392 comparisons out of 12,720) (Fig. S2).

Using a CT threshold of 60, 84 strains were allocated to 21 CTs while 76 strains were reported as singletons (Fig. S3). As previously described (11), 31 of 32 *netF* positive strains clustered into two clusters, CT 1 (*n=*26 strains) and CT 2 (*n=*5 strains). The remaining 19 CTs (CTs 3 to 21) included only poultry strains of *C. perfringens* and were characterised by the inclusion of a few numbers of strains (2 to 5 strains per CT) (Fig. 3).

**Figure 3:**
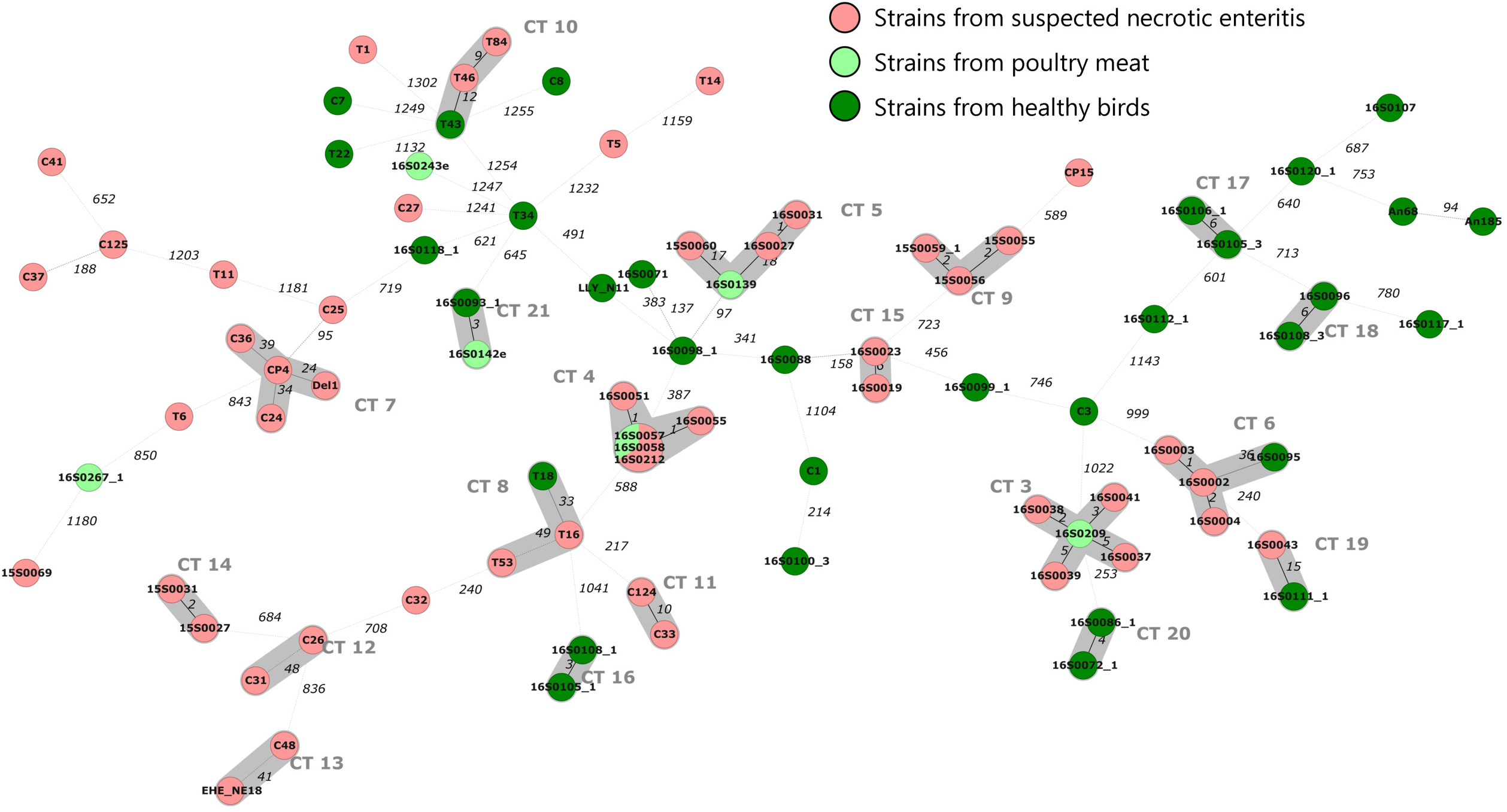
Minimum spanning tree based on the core genome MLST genes of 87 *Clostridium perfringens* strains from poultry, with strains from diseased birds, healthy birds and poultry meat samples highlighted. Circles represent sequence types identified for the strains based on the core genome. Clusters were identified based on fewer than 60 allelic mismatches (grey shading). Numbers on connecting lines illustrate the numbers of target genes with different alleles.

### Genotyping of poultry *C. perfringens* based on cgMLST scheme

The cgMLST scheme was applied to investigate the poultry strains of *C. perfringens*. In total, 87 strains of poultry origin were analysed, 53 thereof were clustered into 19 CTs while 34 strains of poultry origin did not cluster (Fig. 3 and 4). 14 CTs included strains that were isolated from diseased birds. However, seven CTs of these also contained strains from healthy birds and food isolates (Fig. 3).

**Figure 4:**
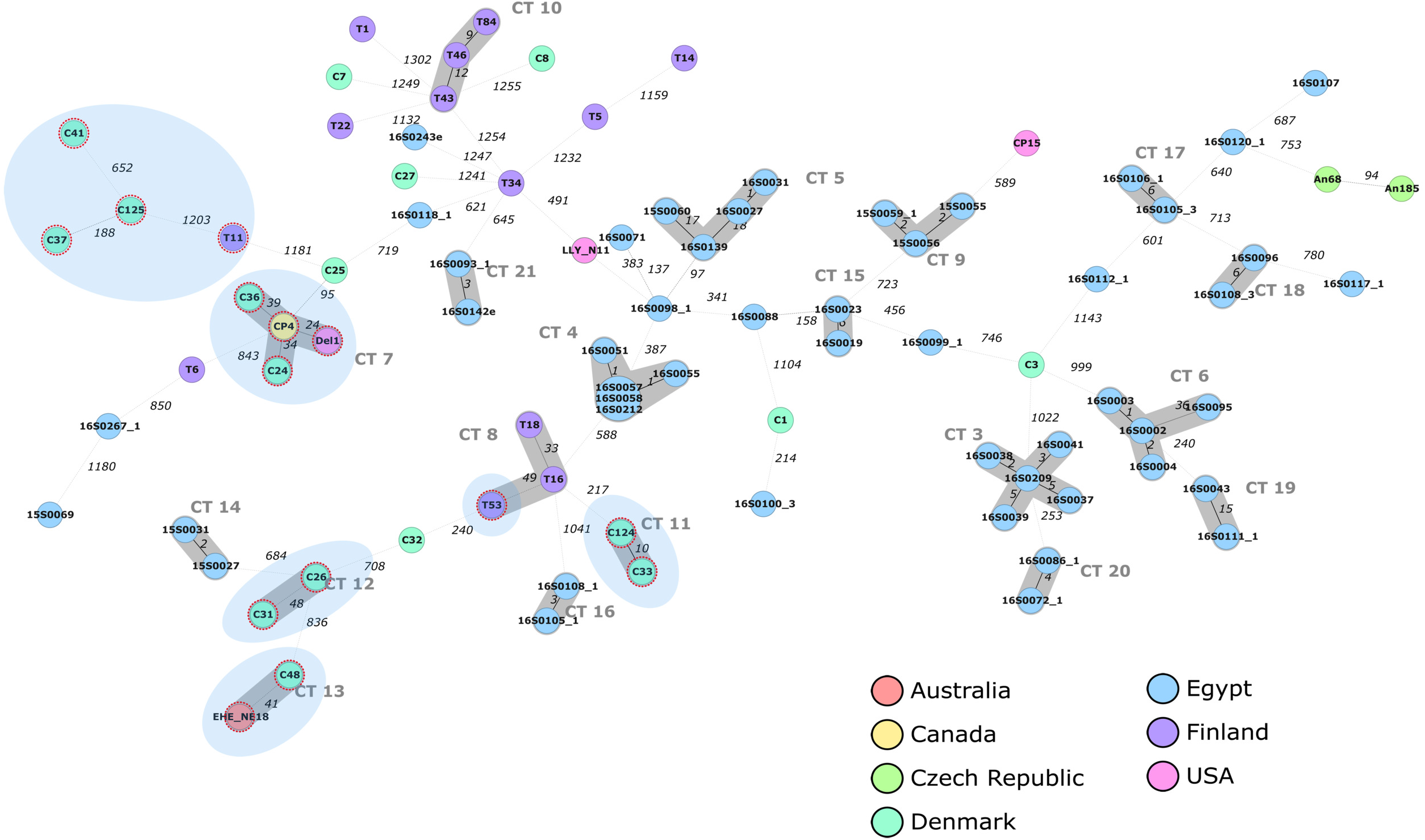
Minimum spanning tree based on the core genome MLST genes of 87 *Clostridium perfringens* strains from poultry with highlighted strains based on the country of isolation. Circles represent cgMLST profiles, and clusters based on fewer than 60 allelic mismatches were indicated with grey shading. The 15 *netB*-positive strains were highlighted with dotted red circles and a light blue shaded background. Numbers on connecting lines illustrate the numbers of target genes with different alleles.

First, cgMLST was applied to explore the relatedness of strains compared to their country of isolation. The results showed that most Egyptian strains grouped into 13 CTs, none of them grouped with strains from the other countries (Fig. 4). Similarly, CT 8 and CT 10 involved only strains from Finland while CT 11 and CT 12 comprised only strains from Denmark. However, CTs including strains from several countries were also detected. For example, CT 7 included four strains from Canada, Denmark and the USA while CT 13 contained two strain from Denmark and Australia (Fig. 4).

Second, cgMLST was applied to investigate the relatedness of 15 *netB*-positive strains (Fig. 4). Ten out of the 15 strains were clustered into four different CTs (CT 7, 11, 12, 13) (Fig. 4). Beside these 10 strains, four additional *netB*-positive strains were identified as phylogenetically related. However, the neighbour allelic distance between them was too high as depicted in Fig. 4.

Lastly, the cgMLST scheme was applied to investigate the 50 poultry strains from Egypt based on the clinical disease and the isolation source, as detailed epidemiological information was only available for these strains (Fig. 5). The results showed that strains originating from the same farm assigned to the same CT. This includes strains from farms C, G, L and N that were clustered and assigned to CT 9, 6, 3 and 4, respectively (Fig. 5). However, two CTs were observed for the strains in farm M, CT 7 (16S0051 and 16S0055) and CT 19 (16S0043). Moreover, despite the strong clustering of the suspected NE isolates in each farm, strains from different farms were sometimes clustered together. For example, CT 4 included strains from farms M and N, CT 9 included three strains from farms C and D, CT 14 included strains from farms A and B, CT 15 included two strains from farm H and I and CT 5 included three strains from farms J, K and E (Fig. 5). Within clusters that include mainly suspected NE isolates, also isolates from healthy birds or poultry meat samples were observed. For example, CT 3, CT 4 and CT 5 included meat isolates and suspected NE isolates. In addition, CT 6 and CT 19 comprise isolates from suspected NE cases as well as isolates from healthy birds.

**Figure 5:**
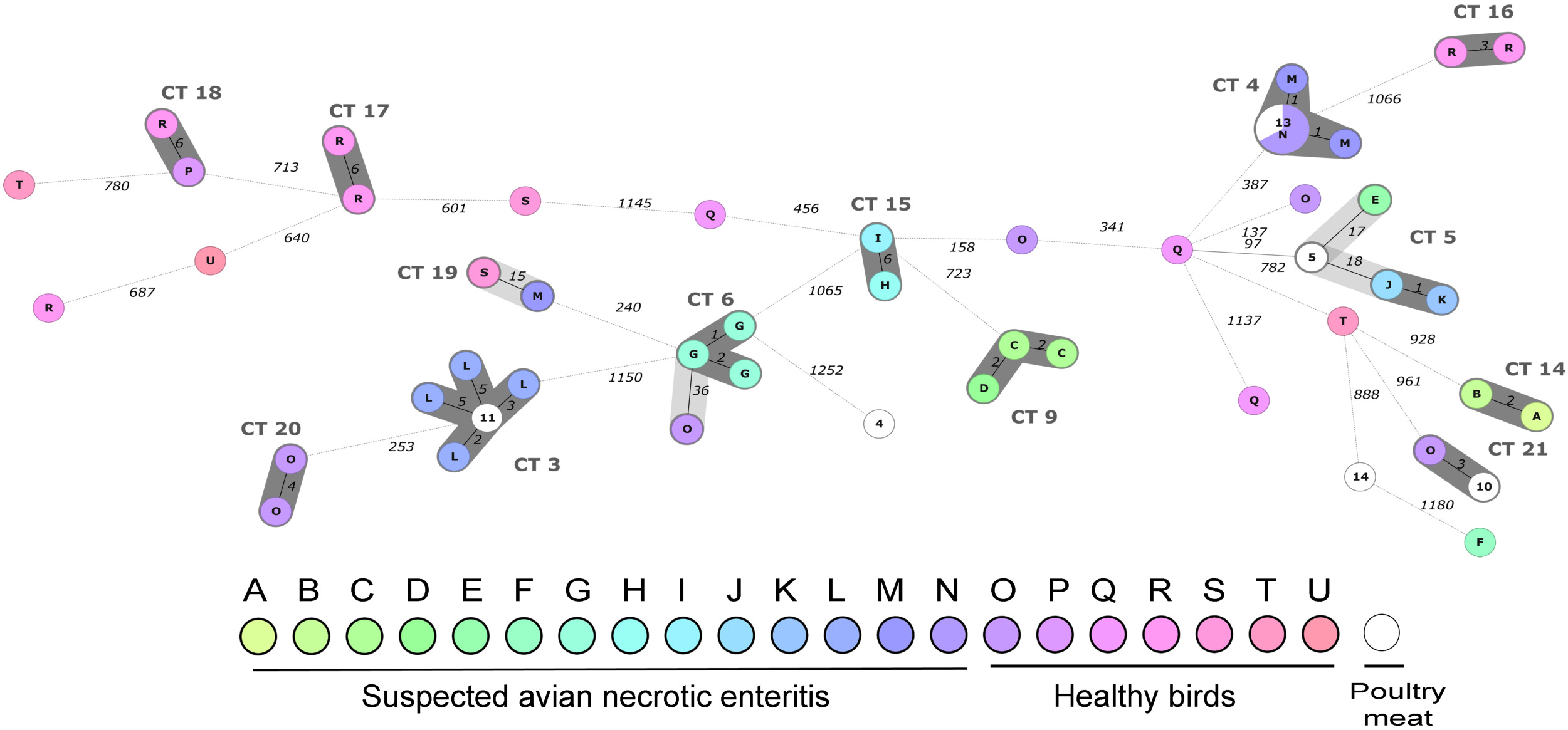
Minimum spanning tree based on the core genome MLST genes of 50 *Clostridium perfringens* strains isolated from poultry in Egypt. Processors A to N refer to the sampled farms in which *C. perfringens* strains were isolated from cases suspected for necrotic enteritis disease. Processors O to U refer to slaughterhouses from which the *C. perfringens* strains were isolated from healthy birds. The nodes were labelled according to the processor name. Nodes labelled with numbers include isolates from poultry meat products. Clusters based on less than 60 allelic mismatches were shaded. The dark shading of the clusters indicates less than ten allelic differences. Numbers on connecting lines illustrate the numbers of target genes with different alleles.

In comparison, *C. perfringens* strains from healthy birds were more diverse even within a single site (Fig. 5). Some strains formed few clusters with a maximum of two strains per cluster as observed for CT 16, CT 17, CT 20 and CT 18 (Fig. 5). Except for CT 18 where the *C. perfringens* strains were derived from two different sites, *C. perfringens* within each of the aforementioned CTs are from a single site.

It is noteworthy to mention that defining the groups based on less than 10 allelic differences maximised the cluster resolution for the epidemiological investigation of the 50 strains from Egypt as it has been shown that the suspected NE isolates did not cluster with any of the non-disease isolates except for two strains in CT 3 and CT 4. Based on less than 10 allelic differences, seven clusters that included suspected NE isolates and four clusters that included non-clinical isolates were found (Fig. 5).

### Comparison between cgMLST cluster analysis and SNP-based phylogeny

Using the tanglegram algorithm (25) there was a high degree of topological congruence between the cgMLST and SNPs-based trees (Fig. S4), where most connecting lines between the corresponding taxa of both trees were in parallel except for very few crossing lines. The 160 *C. perfringens* strains were divided into 156 unique types by the cgMLST while the SNP-based typing resulted in 160 (recombination-unfiltered) and 156 (recombination-filtered) different types for the strains. This resulted in similar SDI (27) for the cgMLST and the whole-genome SNPs (Table S5). These results indicate that both methods provided a maximum resolution for these strains based on Simpson’s index of diversity (27).

### Comparison of core genome MLST and classical MLSTs

MLST schemes described by Jost et al., (2006b), Hibberd et al., (2011) and Deguchi et al., (2009) resulted in 77 STs (SDI = 0.962, 95%CI = [0.945-0.979]), 88 STs (SDI = 0.967, 95%CI = [0.949-0.984]) and 91 STs (SDI = 0.972, 95%CI = [0.956-0.987]), respectively (Table S5). As expected, sequence types inferred from the classical MLST could be further divided into different cgMLST profiles indicating the higher discrimination of the cgMLST compared to classical methods. The degree of agreements between the different typing methods was described as Adjusted Wallace’s coefficient (Table S5). In addition, tree topologies of classical MLSTs were less concordant with cgMLST despite the high discrimination level of all methods (Fig. S5).

## Discussion

Whole-genome sequencing represents a powerful molecular epidemiological tool for pathogen subtyping and outbreak investigations. In this study, we describe a cgMLST scheme for *C. perfringens*. To provide a stable and robust scheme for future analysis, we used a three step procedure to develop the scheme: 1) filtering out 124 unsuitable genes from the reference strain based on repetition, overlapping, truncation or plasmid homology, 2) selecting 1,510 strictly-present core genes in 38 representative strains from the five different phylogroups of the species, and 3) estimating the typeability of genes in geographically and temporally-divergent *C. perfringens* strains (n=80 strains), with further exclusion of frequently untypeable genes (n=71 genes). Therefore, the final cgMLST scheme included 1,431 genes. Compared to the previous cgMLST by Gohari and colleagues, this scheme included more genes and has been evaluated and applied to a relatively larger number of genomes. Furthermore, the scheme has been made publicly available with the establishment of a nomenclature server on the Ridom website (https://www.cgmlst.org/ncs) which can be used with the Seqsphere program. In addition, the scheme is hosted on the PubMLST website (https://pubmlst.org/) as an open-source platform that provides web-accessible analyses for comparative genomics (36). The publicly accessible cgMLST scheme and the allelic nomenclature server should facilitate inter-laboratory comparison of the MLST typing results.

The cgMLST scheme was applied to study 160 *C. perfringens* genomes. The scheme has greater discriminatory power than classical MLST schemes, with strain typing results comparable to whole-genome SNP typing (Table S5). However, in comparison to SNP methods, cgMLST data are easily portable and accessible using online databases. This enables the establishment of a consistent nomenclature (10) which is an essential pre-requirement for global comparison and rapid outbreak communications (37). Moreover, the cgMLST considers gene variation due to insertion or deletion. The inclusion of indel variations in the SNP typing methods is variable and frequently ignored (38, 39). Interestingly, we observed that 298 (20.8%) out of 1,431 of the cgMLST genes were affected by indel mutations. However, cgMLST does not account for variations outside the codring regions. Because of the differences between allele and SNP-calling methods, a combined application of WGS-based MLST and SNPs can provide maximum discrimination for accurate outbreak investigations (53).

Accuracy in detecting genetic variation in a population sample is fundamental to our understanding of bacterial diversity and evolution (40). However, it can be difficult to distinguish true variants from sequencing, assembly and alignment errors (41). The cgMLST method in SeqSphere and BIGSdb is an assembly based system (42). Therefore, sequencing errors and assembly artefacts can affect the results of cgMLST (41). This was indeed observed for the highly fragmented genomes in our data set. A prior study on *Listeria monocytogenes* reported a high reproducibility of cgMLST with different assemblers when the sequencing depth >40-fold (43). In this study, the quality of assembly (n of contigs and N50 value) was low if the coverage was less than 30-fold e.g. for strains T43 and C3 which lack 71 and 29 genes, respectively. However, few other genomes e.g. T34 and C7 had a higher coverage (>80-fold) but also lack 46 and 42 genes, respectively. Therefore, coverage depth may not be the only parameter to influence the results of cgMLST. Other parameters might include the preparation of the DNA and sequencing libraries. The percentage of typeable cgMLST targets per each investigated genome can therefore be used as reflective of the genome quality.

Studies have reported different CT thresholds for the cgMLST typing of different bacterial species. These include fewer than ten alleles to detect epidemiologically linked strains (44). In this study, 50 strains from Egypt were epidemiologically related by geography and isolation time. The 10 allelic differences provided an optimal resolution as strains from diseased birds clustered with only two of the non-disease strains. de Been et al., 2015 described different CT thresholds for *Enterococcus faecium* (45) i.e. >40 alleles for epidemiologically unrelated strains, <20 alleles for closely related strains and between 21 and 40 alleles for possibly related strains (45). Moura and colleagues (2016) described 150 allelic differences between *Listeria monocytogens* strains that belong to the same sublineages (43). In this study, we used a CT threshold of 60 alleles as the frequency of less than 60 pairwise allelic differences was high (Fig. S2). Setting a CT threshold at 60 enabled the clustering of more than half of the investigated genomes (*n=*84) into CTs (*n=*21) (Fig. 3 and 4).

We applied the cgMLST scheme to investigate poultry strains of *C. perfringens* concerning country of isolation, NetB toxin gene carriage and clinical disease. The cgMLST results enabled the identification of CTs that comprise strains derived from a single country (Fig. 4). However, we also identified two CTs that included strains from different countries, and carry the gene of the NetB toxin (Fig. 4), a finding that may indicate common sources of *netB*-positive strains in different countries. This could be due to the poultry commercial system worldwide that is likely maintained by few companies only (46).

The application of cgMLST to 50 poultry strains from Egypt showed that the disease strains recovered from a single farm belong to the same CT. However, strains of the same CT could be detected in different farms. Few CTs could be identified for the suspected NE strains (Fig. 5). This is in contrast to isolates from healthy birds and meat samples. They have a high variety of CTs and most isolates represent singletons. Additionally, suspected NE isolates clustered together with isolates from healthy birds or meat samples. These results are in agreement with a recent genomic study with a focus on NE strains (46), in which the authors reported similar genomic features for NE strains from different geographic regions at various times. This study also showed that the pathogenic clades of NE isolates could not be identified based on the core genome SNP method (46). Taken together, these results show that the disease-causing *C. perfringens* strains within a single farm are usually similar and unrelated strains derived from different suspected NE cases, healthy birds or meat samples could also sometimes cluster together.

## Acknowledgments

We thank Sandra Hennig and Renate Danner for their excellent technical assistance. Mostafa Y. Abdel-Glil received a PhD scholarship from the German Academic Exchange Service (DAAD) within the German Egyptian Research Long-Term Scholarship Program (GERLS).

MA and CS conceived the study. MA designed the study, performed the analysis and wrote the manuscript. PT and JL provided support for data analysis and interpretation. DH and KJ contributed to scheme design and validated the scheme for public release at the Ridom website and PubMLST.org, respectively. LW, HN and CS supervised the study and critically revised the manuscript. All authors approved the final manuscript for publication.

